# Predicting peptide aggregation with protein language model embeddings

**DOI:** 10.1101/2025.09.26.678773

**Authors:** Ethan Eschbach, Kristine Deibler, Deepa Korani, Sebastian Swanson

## Abstract

Amyloid fibrils, a form of peptide aggregate, are associated with multiple diseases and hinder the development of therapeutics. The experimental characterization of aggregating peptides is resource-intensive and data are scarce, limiting the development of accurate models. We present a deep-learning model, PALM (Predicting Aggregation with Language Model embeddings), which uses transfer learning to predict aggregation from embeddings extracted from a pretrained protein language model (pLM). PALM is trained on the WaltzDB-2.0 dataset to classify peptides and identify aggregation-prone regions within a sequence at single-residue resolution. Compared to existing models, it exhibits competitive performance on diverse held-out experimental datasets. We find that PALM fails to identify single mutations that increase the rate of aggregation of amyloid beta peptide; however, training the PALM architecture on a larger dataset, CANYA NNK1-3, substantially improves performance in this task. These results show that transfer learning with pLM embeddings improves performance when training on small datasets, but highlight that challenging tasks, such as predicting the effect of single mutations, require more experimental data.

## Introduction

Amyloid fibrils are filamentous protein aggregates, characterized by their cross-β structure (1). Fibrils form in a sequence-dependent manner, often irreversibly, and have a wide range of biological effects (2). The formation of amyloid fibrils from native human proteins is associated with Alzheimer’s disease, type-2 diabetes, and many other diseases (3). Fibrils pose a challenge to the development of biologic drugs by altering their physical properties and pharmacodynamics (4). Fibrils can also play a functional role, enabling the controlled release of biologically active peptides and displaying unique material properties (5, 6). These challenges and applications motivate the development of models that capture the relationship between amino acid sequence and aggregation.

Many sequence-based methods have been developed to predict aggregation. Most methods use simple descriptors such as hydrophobicity, β-sheet propensity (7, 8), or cross-β pairing energy (9). TANGO—an existing aggregation predictor—uses statistical mechanics to calculate a partition function over secondary structure states (10). While these methods have some predictive ability, they are not machine learning models and cannot be easily improved with new data. These methods are fundamentally limited, as they predict values that are proxies of aggregation. More recently, multiple sequence-based machine learning methods have been developed that learn from experimental data by estimating a position-specific scoring matrix (11), or directly through supervised learning (12, 13). Many of these models have been trained on WaltzDB-2.0, a database of 1,416 hexapeptides classified as amyloid/non-amyloid by Thioflavin-T fluorescence (ThT) assays and Fourier transform infrared spectroscopy (FTIR) (14). Despite the quality of this dataset, it describes only a very small fraction of the potential sequence space of therapeutic peptides.

One strategy to bridge the gap in aggregation data is to leverage models pretrained on large numbers of protein sequences through transfer learning. Protein language models (pLMs) are transformer architectures trained on a self-supervised masked language modeling (MLM) task (15). The representations learned by pLMs have been used for many tasks, including structure prediction, mutational variant effect prediction, peptide properties, and the design of novel functional proteins (16–21). Protein language models have been applied to predict the aggregation of hexapeptides through fine tuning on labelled data, but this work has not yet been extended to peptide sequence fragments longer than six amino acids (22).

We developed a deep-learning model, PALM (Predicting Aggregation with Language Model embeddings), to predict amyloid-forming peptide sequences (Fig.1). PALM is trained on WaltzDB-2.0 and uses embeddings from the pretrained ESM2 protein language model to predict aggregation. PALM extracts local sequence patterns with an adapted Light Attention architecture (23), which we refer to as the Aggregation Predictor Module (APM). The APM infers residue-specific contributions from experimental data and uses them to predict the aggregation of protein sequences with a soft-max weighted mean. To improve prediction for peptide sequences that are longer than the hexapeptides in WaltzDB-2.0, we employ a data augmentation strategy, where non-hydrophobic residues are padded around the original six amino acid WaltzDB sequences prior to training. Relative to existing models, PALM demonstrates strong performance when benchmarked on datasets Serrano157 (10) and AmyPro22 (24), which assess the ability of the model to classify sequences and identify aggregation-promoting regions (APRs), respectively. We further evaluated PALM on additional experimental data, and found that the model fails to identify aggregation-inducing single amino acid substitutions in amyloid beta peptide, which is known to cause familial Alzheimer’s disease. We demonstrate that training PALM on a larger dataset, NNK1-3, improves the performance on this task, as well as sequence-level classification (25).

**Fig. 1.**
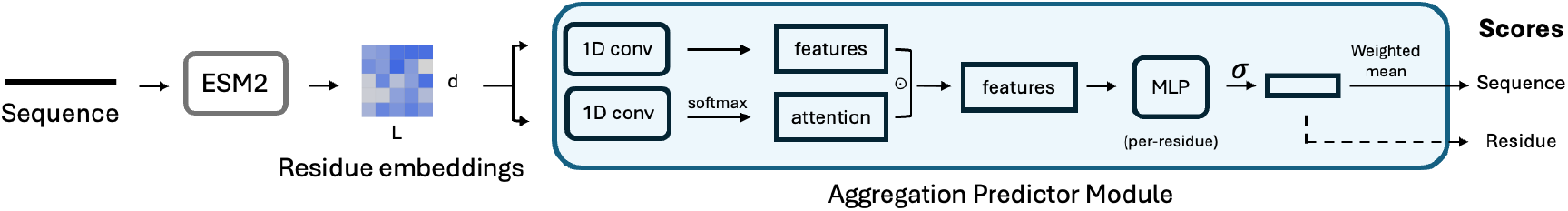
The architecture of the PALM model. Protein sequences are embedded using ESM2 (8M) and then passed to the Aggregation Predictor Module (APM). The module uses one-dimensional convolutions to extract local patterns in the sequence and then predicts the residue score with an MLP. The weighted mean of the residue scores is used to compute the overall score for the sequence. Only the sequence-level score is used to compute the loss during model training.

## Methods

### Datasets

#### Training data

PALM was trained on WaltzDB-2.0, a database of peptides experimentally labeled as amyloid or non-amyloid (14). The database contains 1,416 six amino acid peptides, 515 of which have been observed to form amyloid fibrils. The labels were experimentally determined through ThT assays and FTIR. Prior to model training, the sequences were randomly assigned into five equally sized folds with similar class balances.

The WaltzDB dataset was augmented to diversify the lengths of the sequences in the training data. We modified each six amino acid sequence by appending padding sequence to the beginning and end of the string. The padding strategy implicitly assumes that 1) a six-residue aggregation prone region within a larger sequence is long enough to cause aggregation of the entire protein and 2) that the flanking padding residues would not introduce new APRs into the sequence. The first assumption is shared by other methods that search for short windows within a larger sequence context (10, 12, 13), and is supported by experimental results that show that non-aggregating proteins can be made to aggregate by inserting APRs into their sequence (26, 27). The second assumption depends on the distribution from which the padding residues are sampled, which determines the rate at which new APRs are introduced into the sequences. We specifically sampled non-hydrophobic amino acids to limit the rate at which new APRs were introduced in the flanking region.

The sequences were padded by sampling residues and appending them to both the N and C terminus of the WaltzDB hexapeptide motifs. The length of the padding sequence at each end, *L* was sampled uniformly from the range [1, *L*_*max*_]. As padding residues were added to each end, the maximum possible length of a padded sequence was 2*L*_*max*_ + 6. Three residue type probability distributions were considered: “Random” (uniform), “Mask”, “WaltzDB non-amyloid”, and “Non-hydrophobic”. Random padding was generated by uniformly sampling from the 20 canonical amino acids. All WaltzDB sequences labeled as non-amyloidogenic were used to generate a WaltzDB non-amyloid residue probability distribution. The non-hydrophobic distribution consisted of residues = (C, D, E, G, H, K, N, P, Q, R, S, T), each sampled with equal probability (Fig. S1). To prevent the model from learning spurious patterns between the padding sequence and the APRs, every sequence in WaltzDB-2.0 was oversampled tenfold. Each copy of the repeated hexapeptide was concatenated with unique padding sequences on both ends. (Fig. S2). The oversampling expanded the total size of each augmented dataset to 14,148 sequences and diversified the context in which each hexapeptide was observed. To prevent leakage, the fold assignment of the original six amino acid peptides were preserved with the padded sequences.

#### Evaluation data

Multiple datasets with sequence or residue level binary amyloid labels were used to evaluate the models, as summarized in Table S1. Experimental methods for characterizing amyloid fibril formation are diverse, and sometimes non-amyloid forms of aggregation are unknowingly included in experimental datasets, which can introduce noise in benchmarks. To counteract this, we gathered a variety of datasets annotated through different methods in order to assess our model as thoroughly as possible.

The first dataset, AmyPro, consists of 162 proteins with experimentally-determined residue-level annotation of amyloid-prone regions (24). To prevent data leakage, we filtered out 130 proteins containing hexapeptides found in WaltzDB-2.0 training samples. Following Planas-Iglesias et al. (13), we removed an additional 10 proteins with >50 amino acid amyloid-prone regions, resulting in 22 sequences, which we refer to as AmyPro22. The second dataset consisted of peptides with sequence-level amyloid labels, originally curated by Fernandez-Escamilla et al. to evaluate TANGO (10). The peptides were derived from a total of 24 proteins from multiple publications and measured by CD spectroscopy. In total, this yielded 224 sequences classified as amyloid or non-amyloid. To reduce redundancy to the Waltz training data, sequences were divided into sliding windows of six amino acids and all sequences containing a hexapeptide motif found in WaltzDB were removed. To minimize overlap between evaluation sets, we removed two sequences with greater than 40% sequence similarity to sequences in AmyPro22, as determined by global sequence alignments with the Needleman-Wunsch algorithm (28). We removed a single duplicate of peptide m68 (29), leaving only the positive label. This left a total of 157 sequences, which we refer to as Serrano157.

Concurrently with this work, a large dataset of >100,000 peptides tested for aggregation via a massively parallel selection assay became available (25). The sequences are between 1 and 20 amino acids, and were obtained using synthetic NNK oligo pools, resulting in randomized peptides with a high degree of sequence diversity. An aggregation score is reported for each sequence, as determined by an *in-vivo* selection assay with binary labels assigned by the authors. The test set, NNK4, contains 7040 sequences. The training set, NNK1-3, contains 100,730 sequences. The sequences were randomly assigned to five folds for cross-validation.

In an earlier work, a similar selection assay was used to measure aggregation scores for mutants of Aβ42 peptide (30). Of the 753 single amino acid substitutions in this dataset, 13 are recognized to cause familial Alzheimer’s disease by driving amyloid formation (31, 32). Following Seuma et al., we define the 13 fAD substitutions as amyloid and the remaining substitutions as non-amyloid. We refer to this dataset as Aβ42 peptide mutants.

### Protein language model embeddings

We used the pretrained protein language model, ESM2, to extract embeddings from amino acid sequences (16). ESM2 is a transformer trained with a masked-language modeling (MLM) objective on sequences from UniRef50. To explore the influence of scale on the embeddings, we considered four model sizes: 8M parameters (6 layers), 35M parameters (12 layers), 150M parameters (30 layers), and 650M parameters (33 layers). For each sequence, an embedding tensor of shape [*L, d*_in_], where *L* is the length of the sequence and *d*_in_ is the size of the hidden dimension, was extracted from the final hidden layer of the model. The value of *d*_in_ varies depending on the size of the model. To visualize the embeddings, the embedding vectors were mean-pooled along the length dimension and dimensionality reduction was performed with t-stochastic neighbor embedding (t-SNE). For the feature ablations, we used 5-dimensional z-scales derived from physicochemical descriptors (33).

### Aggregation Predictor Module architecture

The Aggregation Predictor Module (APM) was designed to identify local sequence motifs that form amyloid-promoting regions. The APM was adapted from Light Attention (23), which uses convolutional filters to extract local patterns in the residue embeddings generated by pLMs. As described previously, embeddings with dimensions [*L, d*_*in*_], were extracted from ESM2 and passed to the APM. Then two independent, one-dimensional convolutions (kernel size = 5 and stride = 1) were applied along the sequence dimension to yield a value tensor **v** and attention tensor **a**, both with dimensions [*L, d*_*in*_] (the hidden layer dimension size of the original embeddings was preserved). Softmax was applied over the *L* dimension of the attention tensor to convert the values to attention weights. Dropout was applied with a probability of 0.25 to the feature tensor. The two tensors were then multiplied element-wise to generate a new attention-weighted feature tensor **f** with dimensions [*L, d*_*in*_]. The tensor **f** was passed to a multi-layer perceptron (MLP), which operates along the *L* dimension. The MLP consisted of two linear layers (output dim=32, output dim=1). Dropout (rate = 0.25), ReLU activation, and batch normalization were applied between these layers. The output was transformed with a sigmoid function to yield a vector **r** of length *L* with values in the range [0, 1]. The values in **r** are residue importance scores inferred by the model (which we refer to as residue scores for brevity). The residues scores were used to compute the sequence score *s* by applying a weighted mean operation

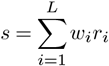

Where *w*_*i*_ is the residue-specific weight computed from **r** by applying softmax over the sequence length. The residue-wise application of the MLP and the weighted mean are the main modifications to the original Light Attention module. The use of the weighted mean to compute *s* from the values in **r** enhances the interpretability of the model.

### Model training

The model was trained to minimize the binary cross-entropy loss function using stochastic gradient descent. The loss was averaged for each batch (n=1000). For a predicted sequence score 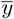, the loss was defined as:

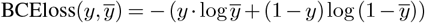

To avoid overfitting, the model was trained with early stopping and checkpointing. If the validation loss did not decrease after 200 epochs, the weights from the epoch with the lowest validation loss were selected. Hyperparameter optimization was performed through a manual grid search. Convolutional kernel size (1, 5, 7, 11), learning rates = (0.5, 0.05), and MLP layer count (2, 4) were explored. Hyperparameters ranges were limited to small values to limit the total number of parameters in the model. The final hyperparameter values were selected based on 5-fold cross validation loss on WaltzDB. The final hyperparameter values that were selected were kernel size = 5, learning rate = 0.05, and MLP layer count = 2.

### Model evaluation

We used five-fold cross-validation to evaluate the models. We computed ROC AUC, AUPRC, and the enrichment factor, i.e, the fold enrichment in precision at top 10%. We considered four evaluation datasets: Aβ42 mutants, NNK4, Serrano157, and AmyPro22. For the datasets annotated at the sequence level (Serrano157, Aβ42 mutants, NNK4), each sequence was taken to be an individual sample. For residue-level annotations (AmyPro22), individual amino acids were considered to be a single sample.

Serrano157 and AmyPro22 were used to perform ablation studies on PALM. Ablations applied to the main version of PALM were compared using Tukey’s Honestly Significant Difference (HSD) test computed using the statsmodels python package (34). The model ablations were grouped into three categories: dataset, embedding, and architecture. HSD tests were performed independently on each group of ablations.

We evaluated multiple versions of PALM, with different residue padding strategies and ESM2 model sizes to understand their importance and select the optimal configuration. For these experiments, we used fixed hyperparameters identified during the manual grid search. We selected the non-hydrophobic, *L*_*max*_ = 10 residue padding strategy and ESM2 8M based on the performance of the model on Serrano157 and AmyPro22. We also performed ablations on the APM, removing sections of the module and retraining the model. For a complete ablation, we replaced the entire APM with a logistic regression (LR) head. The LR head was trained using nested cross-validation over the five folds. The hyperparameters were optimized with grid search in the inner loop of the cross-validation. We explored penalties: (l1, l2) and regularization strength: (0.01, 0.1, 1.0, 10, 100). The comparisons between padding strategies, ESM2 model sizes, and ablations are shown in Figure 3.

We benchmarked PALM against TANGO (10), AggreProt (13), AggreScan (35), ANuPP (12), CANYA (25), PASTA (36) and Waltz (11). With the exception of CANYA and Waltz, these methods output residue-level scores, which we used to compute performance metrics on AmyPro22. We considered sequences to be classified by Aggrescan as amyloid if the number of hotspots was one or greater. For models that only provide residue values, including PASTA, ANUPP and AggreProt, we obtained sequence-level values by taking the maximum over the residue values. In all cases, taking the maximum residue probability as the overall sequence probability outperformed the mean.

We addressed hydrophobicity biases on sequence and residue score assignment by splitting each validation dataset into low and high hydrophobicity groups. For Serrano157 and CANYA NNK4, the per-residue Kyte-Doolittle hydrophobicity index was averaged across each sequence. This average hydrophobicity value was used to stratify each dataset into two even-sized groups. For AmyPro22, the average Kyte-Doolittle score (37) for each sequence was calculated using only the non-APR regions of the sequence. The 11 sequences with the highest average background Kyte-Doolittle scores were assigned to the high hydrophobicity group, while the 11 lowest were assigned to the low hydrophobicity group.

## Results

### Adding padding residues to WaltzDB sequences aligns their ESM2 representations with Serrano157 and AmyPro22

The hexapeptides in WaltzDB are shorter than the majority of natural peptides and the sequences in Serrano157 and AmyPro22 (Fig. 2). We visualized the embedding space of ESM2 8M using t-SNE and observed that the sequences in WaltzDB form a distinct cluster that does not overlap with the sequences in Serrano157 and AmyPro22 (Fig. 2). Notably, the sequences did not cluster by amyloid classification, indicating that local sequence features, not global embedding similarity, were more responsible for those differences. We hypothesized that the difference in sequence length was partially responsible for the observed clustering, indicating that adding additional residues to the WaltzDB sequences could counteract this length-dependent global difference between the hexapeptides and longer sequences. Each sequence in WaltzDB was modified by adding up to 10 additional randomly sampled residues per terminal end to introduce variation in the total length of the sequence (see methods). The sequences in WaltzDB-padded (non-hydrophobic, *L*_*max*_ = 10) formed a cluster that overlapped the Serrano157 cluster, distant from the original WaltzDB cluster. Other padding strategies, which varied in the amino acid composition and maximum length of the padding region, exhibited similar trends (Fig. S3).

**Fig. 2.**
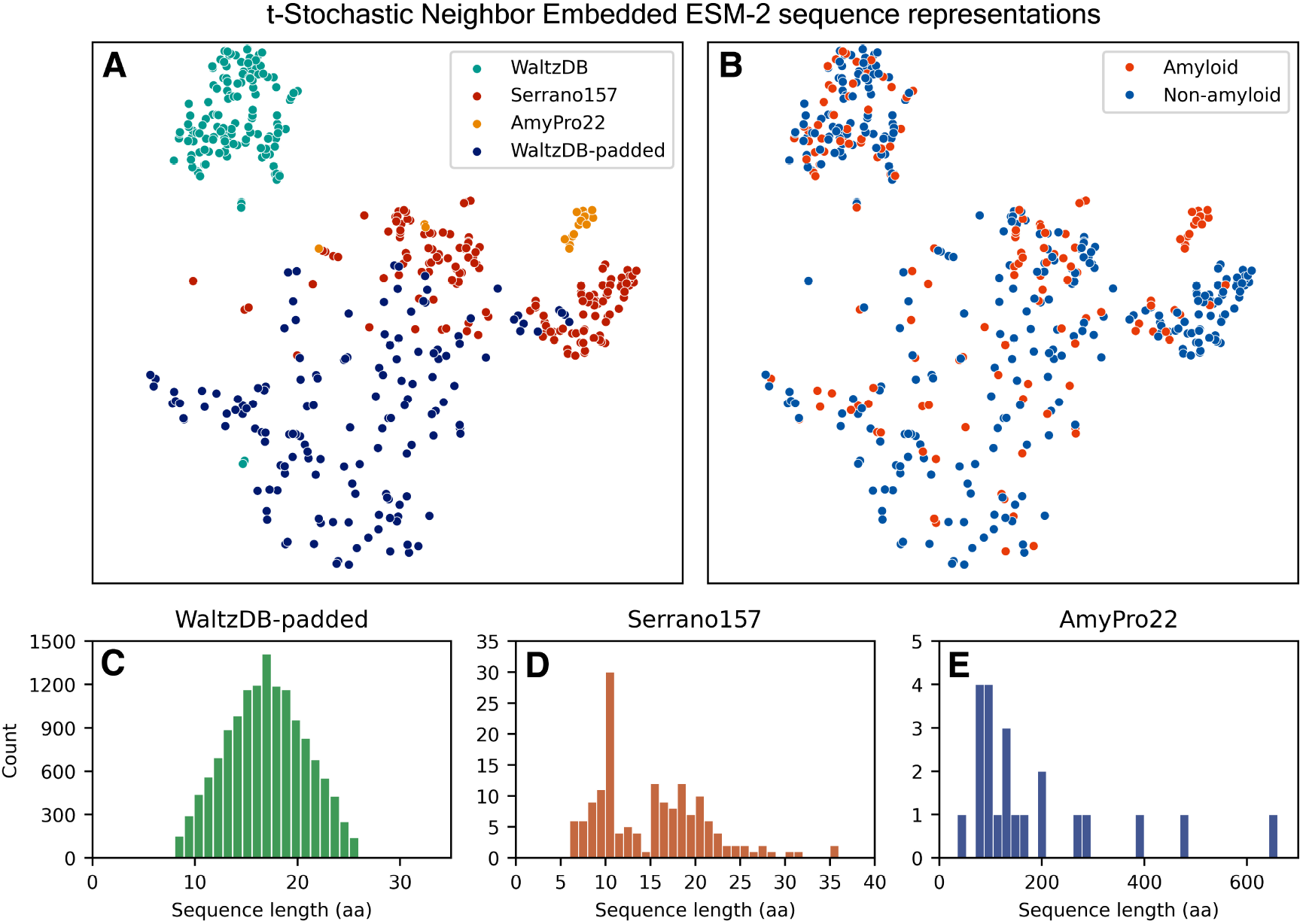
The sequences in WaltzDB form a distinct cluster from other peptide sequences. The ESM2 8M embeddings for sequences from WaltzDB, Serrano157, AmyPro22, and WaltzDB-padded (non-hydrophobic, *Lmax* = 10) are visualized using t-SNE dimensionality reduction. The sequences are colored by dataset (A) and by their Amyloid/Non-amyloid class label (B). The WaltzDB and WaltzDB-padded (non-hydrophobic, *Lmax* = 10) datasets were sub-sampled to 200 sequences for visualization. The distribution of sequence lengths for WaltzDB-padded (C), Serrano157 (D), and AmyPro22 (E).

### WaltzDB residue padding improves sequence and residue classification performance

We trained PALM with WaltzDB padded using different strategies (see methods) and measured sequence classification performance on Serrano157 and residue classification on AmyPro22. The highest performance was observed with WaltzDB-padded (Non-hydrophobic, *L*_*max*_ = 10aa), which significantly improved performance on both datasets relative to the model trained on unaltered WaltzDB sequences (Fig. 3). The difference in ROC AUC was more pronounced on Serrano157, suggesting that padding the training data was particularly important for sequence-level classification. Alternative residue padding strategies performed significantly worse on one or both evaluation datasets, with the exception of non-hydrophobic amino acids with a maximum length of 20 aa. This strategy appeared to be worse than 10 aa non-hydrophobic padding; however, HSD tests indicated that this performance difference was not statistically significant. We inspected the residue level scores of the sequences in the validation set and found that the models trained with alternative padding strategies assigned higher scores to residues from the non-amyloid WaltzDB sequences and padding regions (Fig. S4). The median residue score was found to be higher in the padding and non-amyloid residues for models trained with all other padding strategies, potentially because they erroneously introduce APRs into the padding region at a higher rate. Across strategies, the median residue score of the padding and non-amyloid residue scores were similar, supporting that there is not a strong systematic bias of disproportionately low scores for the padding positions learned during training.

**Fig. 3.**
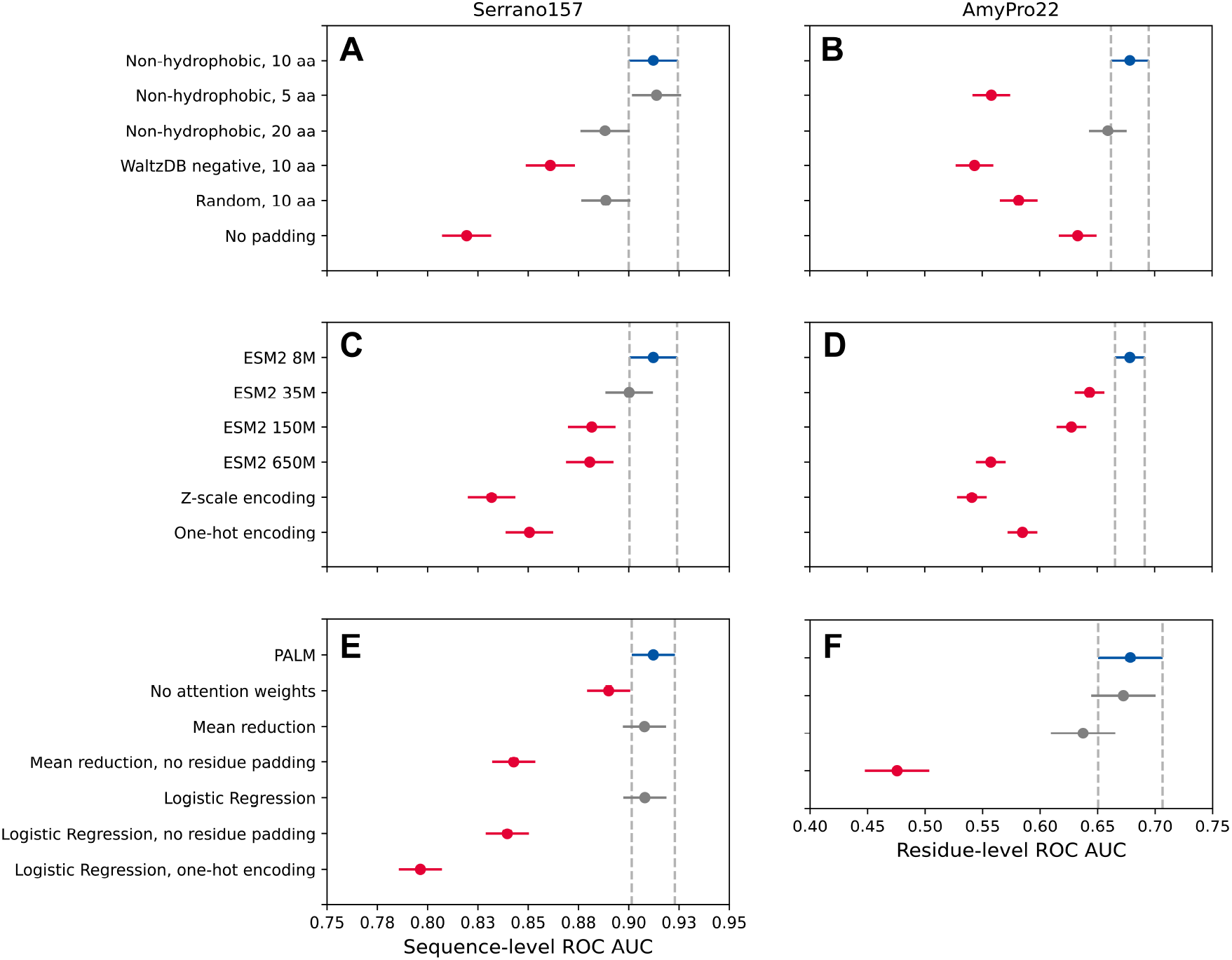
Ablations reveal key elements of the model. ROC AUC scores across cross-validation folds were compared using Tukey’s Honestly Significant Difference (HSD) test. PALM is shown in blue with a dashed confidence interval surrounding it. Model ablations that performed comparatively to PALM are shown in grey, and models that fall outside of the confidence interval are shown in red. Results are shown for Serrano157 (A, C, E) and AmyPro22 (B, D, F) datasets, comparing residue padding distributions and maximum padding lengths per terminus (A, B), embedding models (C, D), and model architectures (E, F).

### The smallest ESM2 model provides the best performance

To explore the influence of language model scale on amyloid prediction, we trained the APM using four ESM2 models with increasing numbers of parameters. We observed a negative relationship between the size of the ESM2 model and the performance on both test datasets. Substituting ESM2 embeddings with one-hot or z-scale encodings significantly lowered performance on Serrano157 and AmyPro22, supporting that embeddings from the protein language model substantially contribute to the models predictions (Fig. 3).

The Serrano157 ROC AUC taken from the ESM2 8M model (PALM) was 0.91 ± 0.005 was significantly better than all other sizes, with the exception of the 35M parameter model. The AmyPro22 ROC AUC (0.68 ± 0.02) was significantly higher than all other models. In general, the drop in performance with larger language models (ESM2 650M) relative to the default (ESM2 8M) was more pronounced with the AmyPro22 APR prediction task. Between the largest and smallest embedders tested, the ROC AUC dropped by 0.120 on AmyPro22 and 0.031 on Serrano157. We observed that the loss at the final epoch consistently decreased with larger model sizes, while the validation loss increased, suggesting some degree of overfitting when PALM was trained with features from the larger ESM2 models (Table S2).

### The APM enables residue-level prediction of aggregation-promoting regions

We found that replacing the light attention module with a simple one-dimensional CNN (ablating the attention tensor) resulted in a small, marginally significant drop in performance on Serrano157 (Fig. 3). This change did not impact residue-level prediction performance. Similarly, replacing the weighted-mean reduction operation with the mean over the residue scores did not significantly impact performance on either task. However, when the model was trained with mean-reduction and WaltzDB (no padding), the ROC AUC on AmyPro22 dropped substantially. This suggests that the weighted mean is beneficial for the detection of APRs in sequences with highly variable lengths. Surprisingly, replacing the predictor head with a simple logistic regression (LR) model resulted in equivalent performance on Serrano157. Removing padding and featurizing the sequences with one-hot encodings led to a large decrease in ROC AUC. The drop in ROC AUC observed with the use of one-hot encodings was more pronounced when paired with logistic regression than it was with the APM.

### PALM predicts aggregating peptides and aggregation-prone regions

#### Comparison to existing models

We measured the performance of our model against established methods for predicting amyloid aggregation using the Serrano157 and AmyPro22 datasets (Table 1). PALM exhibited strong performance on both datasets, with other models performing moderately worse on one or both. PASTA (36) exhibited a remarkably high ROC AUC on AmyPro22, but showed a lower enrichment and AUPRC, and notably failed to predict sequence-level aggregation as assessed by Serrano157. The remaining methods performed similarly on AmyPro22, with ROC AUC values ranging from 0.639 (AggreProt) to 0.649 (ANuPP). There was a wider range of performance on Serrano157 relative to PALM: Aggrescan and Waltz performed notably worse, ANuPP only moderately worse, and TANGO and Aggreprot performed comparably. The AUPRC and enrichment values were highest for PALM on Serrano157, but comparable to existing methods on AmyPro22. Overall, TANGO came the closest to matching the performance of PALM.

**Table 1.** Classification metrics measured on residue-level (AmyPro22) and sequence-level (Serrano157) predictions across multiple models.

| Model | Serrano157 |  |  | AmyPro22 |  |  |
| --- | --- | --- | --- | --- | --- | --- |
|  | ROC AUC | AUPRC | Enrichment factor | ROC AUC | AUPRC | Enrichment factor |
| PALM | $0.918 \pm 0.003$ | $0.770 \pm 0.013$ | $3.65 \pm 0.12$ | $0.678 \pm 0.018$ | $0.360 \pm 0.010$ | $2.27 \pm 0.03$ |
| TANGO | 0.894 | 0.691 | 2.53 | 0.645 | 0.370 | 2.72 |
| AggreProt | 0.888 | 0.689 | 2.79 | 0.639 | 0.291 | 2.19 |
| ANuPP | 0.851 | 0.657 | 3.29 | 0.649 | 0.338 | 2.46 |
| CANYA | 0.822 | 0.599 | 2.53 | N/A | N/A | N/A |
| AggreScan | 0.817 | 0.669 | 2.79 | 0.645 | 0.335 | 2.31 |
| Waltz | 0.794 | 0.462 | 2.03 | N/A | N/A | N/A |
| PASTA | 0.327 | 0.181 | 1.20 | 0.707 | 0.220 | 1.20 |

The effects of sequence length on model performance were further evaluated by stratifying the Serrano157 and AmyPro22 datasets by length (Table S3). For residue-level classification, PALM performance was consistent across both groups of AmyPro sequences. PALM sequence-level classification performance was lower for longer sequences in Serrano157, with ROC AUC values of 0.85 on sequences longer than 18 amino acids and 0.92-0.90 on two groups of shorter sequences. TANGO exhibited similar trends, with ROC AUC values of 0.77 on Serrano157 sequences longer than 18 amino acids and 0.94-0.99 on shorter sequence groups.

#### Sequence and residue score distributions

We observed differences in the distribution of residue and sequence scores when grouping by the label from the evaluation datasets (Fig. 4). The sequence scores on Serrano157 were bimodal. While the majority of the non-amyloid sequences had scores close to 0, some were erroneously assigned scores higher than 0.5. In contrast, almost all of the amyloid sequences were assigned scores close to 1. The residue score distributions on AmyPro22 were unimodal and centered around 0.78. The amyloid sequences exhibited a tail to the right with higher scores. The non-amyloid sequences did as well, but to a lesser degree. In comparison, there was a much higher degree of separation of scores between classes in the training set.

**Fig. 4.**
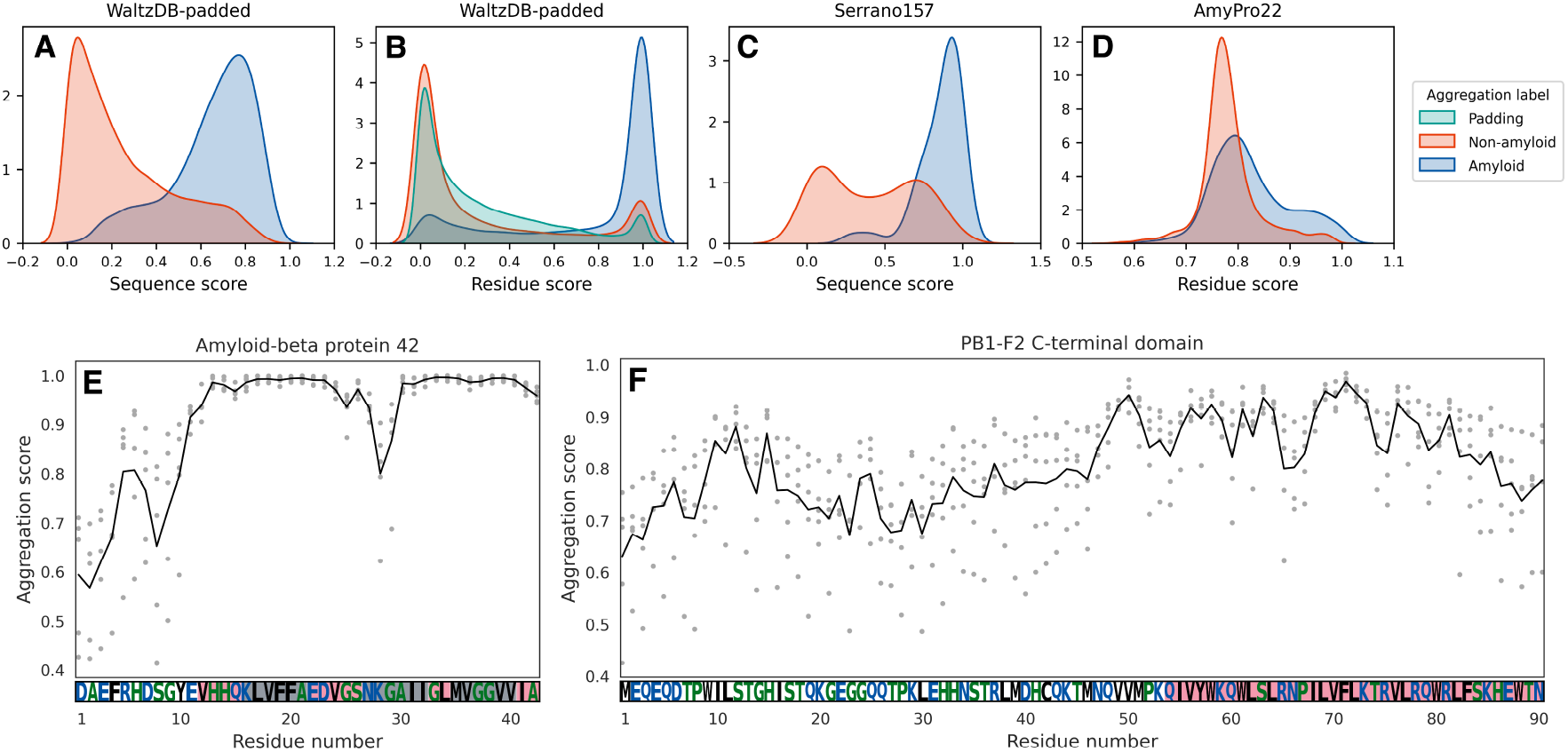
PALM learns to identify aggregation-prone regions in peptide sequences. Top row: comparison of sequence and residue score distributions. The sequence score for amyloid/non-amyloid sequences in the training set (A), the residue score of amyloid/non-amyloid/padding residues in the training set (B), the sequence score for Serrano157 (C), and the residue score for AmyPro22 (D) are shown above. The amyloid/non-amyloid distributions of the test set sequences show separation, with some overlap. Bottom row: residue score profiles for human amyloid beta peptide 42 (E) and PB1-F2 influenza A (F). The APR residue range from AmyPro22 is highlighted in red. The hexapeptides present in the WaltzDB2.0 training set are highlighted in green.

To understand the degree to which the sequences and residue scores were driven by hydrophobicity, we looked at the correlation between the scores and the Kyte-Doolittle hydrophobicity index (Table S4). We found that the scores predicted for Serrano157, AmyPro22, and NNK4 evaluation datasets weakly correlate with hydrophobicity. Spearman r values between residue-level model scores and the hydrophobicity index values did not exceed 0.23. Correlations between sequence-level scores and averaged hydrophobicity index values did not exceed 0.32, with the exception of Serrano157 (Spearman r=0.52); however, the base correlation between average hydrophobicity index for Serrano157 against sequence class labels was already high (Spearman r=0.41).

We also considered whether the non-hydrophobic padding strategy might lead to bias in the scores of certain residues types. We found that the median residue scores of valine, isoleucine, and leucine were elevated relative to the other amino acids in the AmyPro22 dataset (Fig. S5). Interestingly, the other residues excluded from the padding were not the most elevated, and methionine in particular had among the lowest median score of all amino acids (Fig. S1).

#### Visualizing residue score profiles

The residue score profiles of representative peptides highlight aggregation-prone regions within the sequence. First, we considered the 42 amino acid isoform of human amyloid beta peptide (Aβ42), which contains an APR between residues 12-42. The residue scores were highest in two regions between residues 13-23 and 30-40 (Fig. 4E). Both of these regions match sequences labeled as amyloid in WaltzDB. The residue scores were notably high for all five models trained with cross-validation, indicating that the high aggregation scores persist even when those sequences are held out in the validation set. We also considered the viral protein PB1-F2, which contains a C-terminal region (i.e., residues 53-90) that has been shown to form amyloid fibrils (38) (Fig. 4E). The residue score profile of PB1-F2 displayed multiple peaks, with the highest between residues 68-73, part of the region that is annotated. Some residues outside of the C-terminal region also showed higher scores, but overall the values were similar to the average score across the AmyPro22 dataset (0.78). Prediction variability between five crossvalidation folds was generally observed to be at its lowest in amyloid-prone regions. Scores across all five folds exhibited near-perfect agreement in the aggregating regions of the 42-residue amyloid beta peptide. Interestingly, the residue scores were lower between residues 24-29, despite the fact that this contains a third WaltzDB sequence labeled as amyloid, potentially because it plays less of a role in aggregation. Regions of the PB1-F2 APR exhibit a similar lack of variability across cross-validation folds, including residues 68 to 72.

### Evaluating PALM on additional datasets

#### PALM classifies diverse sequences, but struggles to predict the effect of amino acid substitutions

We evaluated PALM on additional datasets to assess its practical utility. First, we considered NNK4, (25) a random library of peptides experimentally labeled as amyloid/non-amyloid by a high-throughput parallel selection assay. PALM showed some predictive power, with a slightly higher ROC AUC than TANGO and AggreProt, but lagged behind CANYA, a model trained on a large dataset obtained with the same high-throughput experimental assay (Table 2).

**Table 2.**
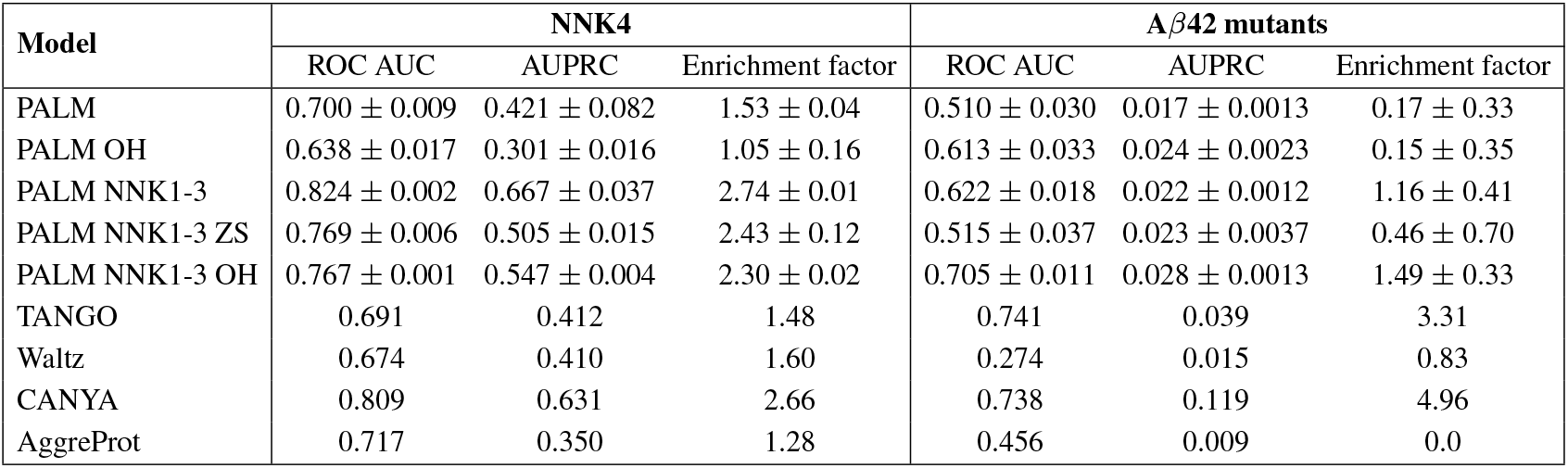
Classification metrics calculated from sequence-level predictions of the NNK4 and A*β*42 datasets across multiple models.

Next, we considered single amino acid substitutions of amyloid beta peptide (Aβ42). We followed Seuma et al. (30), evaluating the ability of the model to assign a higher score to 13 variants of Aβ42 with single amino acid substitutions known to cause familial Alzheimer’s Disease (fAD) by increasing the rate of aggregation. Surprisingly, we found that only TANGO and CANYA performed well on this task, whereas PALM, Waltz, and AggreProt completely failed to identify these mutations (Table 2).

#### Retraining on CANYA data improves performance on NNK4 and Aβ42

We retrained the PALM architecture with the NNK1-3 dataset with features from ESM2 8M (PALM NNK1-3), one-hot encodings (PALM NNK1-3 OH), and z-scales (PALM NNK1-3 ZS). PALM NNK1-3 matched the performance of CANYA on the NNK4 test set and showed an improved ability to predict dominant fAD mutations (Table 2). Interestingly, relative PALM NNK1-3 (ESM2 8M), PALM NNK1-3 OH was less effective on the NNK4 dataset, but showed improved performance on the fAD mutation prediction task. We also benchmarked the PALM ablation trained with one-hot encodings and found that it displayed predictive power on Aβ42, with a substantially higher ROC AUC and AUPRC than base PALM.

We split each diverse validation dataset (Serrano157, AmyPro22, and NNK4) into low- and high-hydrophobicity groups by Kyte-Doolittle indices and repeated the analyses above (Table S5). For residue-level classification (AmyPro22), all models exhibited roughly constant performance across both hydrophobicity groups. Sequence-level classification performance on NNK4 was also roughly constant across both groups for all models. Due to its limited size and strong correlation between class labels and hydrophobicity (Spearman r=0.41), Serrano157 could not be split into two hydrophobicity groups with equal amyloid class label distributions.

We visualized the residue scores profiles of WT Aβ42 and the 13 dominant fAD mutants from PALM and PALM trained on NNK1-3 (Fig. 5). The residue scores predicted by PALM for each of the mutants were similar to the wild-type sequence, which already predicted values close to 1 at residues 12-42, despite the fact that the mutations are known to further increase the rate of aggregation. The residue score profiles predicted by PALM (NNK1-3) were lower overall (Fig. S6), but displayed sharp peaks in similar regions in the sequence (Fig. 5). In particular, Val18 and Phe19 were respectively predicted by PALM (NNK1-3) and PALM (NNK1-3, one-hot) to be among the most aggregation prone residues of the sequence. The difference in residue scores between the mutants and the wild-type predicted by PALM (NNK1-3, one-hot) were localized to within 1-3 residues, as expected by the size of the convolutional filter. In contrast, some single mutations were predicted by PALM (NNK1-3) to have a broader effect. For example, the residue score for the E22G Arctic mutant at position 22 was slightly lower than WT, but increased from residues 25-42. One of the mutations that was not predicted significantly increase aggregation, A21G, was experimentally determined as having the smallest positive effect on aggregation of the 13 mutants (30).

**Fig. 5.**
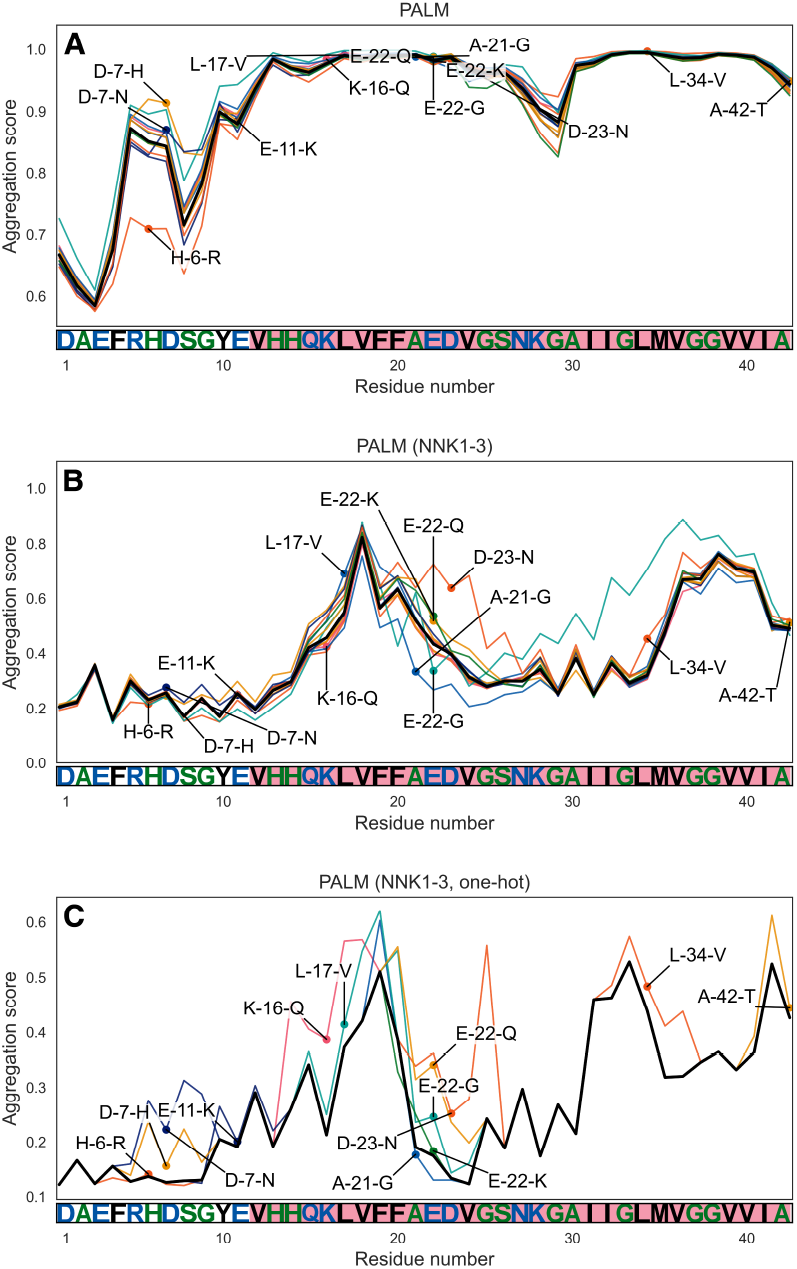
Training PALM with NNK1-3 leads to improved detection of mutations that promote aggregation in familial Alzheimer’s disease. The residue scores for the wild-type Aβ42 sequence are shown as a black line and the scores for the sequences with aggregation promoting substitutions are shown as colored lines. Mutation profiles are included for PALM (A), PALM trained on NNK1-3 (B), and PALM trained on NNK1-3 with one-hot encodings (C). The mutated residue is highlighted with a dot and label. The wild-type sequence and residue number is shown below. The residue range annotated as the APR in AmyPro is highlighted in red.

## Discussion

### PALM classifies peptides using embeddings from ESM2

We present a model trained on WaltzDB-padded that leverages ESM2 8M embeddings to achieve robust prediction of amyloid peptides in the benchmark datasets Serrano157, and NNK4. Replacing the sequence embeddings with one-hot encodings or z-scales results in a large drop in performance, supporting that the embeddings are key to the model’s performance. The observation that z-scales do not improve over one-hot encodings, is surprising, as they contain physico-chemical information. We hypothesize that the embeddings contain additional information beyond residue-level physical features, such as secondary structure propensity and evolutionary constraints on aggregation-prone sequences, which are learned during pretraining. The importance of the ESM2 embeddings is also reflected in the strong performance of the logistic regression ablation on sequence-level classification.

Despite the importance of pLM embeddings, we found that increasing the scale of the ESM2 model leads to decreased performance. This is in contrast to the observed scaling relationships for contact and 3D structure prediction (16). We note that other work has shown the benefit of increased scale is only observed tasks that are not closely aligned with the masked language modeling pretraining task (39). Li et al. do observed a positive scaling relationship between ESM model size and secondary structure prediction, but it is important to note that their evaluation sets consist of natural proteins with well-defined folds, whereas the training sets considered in our work (WaltzDB, WaltzDB-padded, NNK1-3) consist of short peptide sequences. We hypothesize that the larger ESM2 embedding vectors contain additional information, e.g. evolutionary information such as residue conservation due to functional constraints, which is unrelated to the aggregation prediction task. The observed drop in performance may also arise from insufficient regularization in the PALM architecture.

The addition of padding residues is important to the performance of the model on Serrano157, likely because the sequences in the evaluation sets are substantially longer than the six amino acid sequences in WaltzDB-2.0, leading to a shift in the embedding space that is detrimental to model performance. Despite the improved performance, we note the non-hydrophobic padding strategy does not reflect complex phenomena such as long-range cooperative effects, which could be observed in longer sequences. While we cannot rule out subtle biases introduced by the non-hydrophobic amino acid composition of the padding, comparable performance of alternate padding strategies and performance across diverse evaluation datasets supports that PALM has learned to predict from features beyond hydrophobicity.

The performance of PALM on Serrano157 and NNK4 is robust relative to established methods, with only TANGO and Aggreprot close to matching its performance. However, on NNK4 there is still a gap between PALM and CANYA. This is partially explained by the fact that CANYA was trained on data from the same assay. We show that when PALM is trained on the same data (NNK1-3) it achieves similar performance. Altogether, this suggests that the ESM2 8M embeddings are beneficial for training an amyloid sequence classification model, but are not sufficient to compensate for the limited sequence diversity in WaltzDB-2.0, which is 100 times smaller than NNK1-3. The ablations revealed that replacing the APM with logistic regression does not significantly impact the performance on Serrano158, suggesting is not critical for the sequence-level task. Future work can explore modifications to the reduction operation in the APM that allow for the cumulative effect of multiple APRs to be better captured.

### The ability to identify APRs emerges without training on residue-level labels

Identification of APRs within sequences is critical for understanding how individual residues contribute to aggregation. Due to practical considerations and limitations in available data, many works in the field have focused on short windows of longer peptide sequences when predicting APRs (10–12, 22). In contrast, PALM generates embeddings of the complete sequence, allowing for long-range interactions to be captured by the protein language model, which can be extracted by the APM.

PALM performs well relative to al existing models, except for PASTA, when identifying residues within APRs in AmyPro22, and this ability emerges from the model without explicitly being trained to predict aggregation-promoting residues. The shift in residue scores distributions across datasets reflects that the values are best suited for ranking and highlighting potentially problematic positions, as opposed to direct classification.

The performance on AmyPro22 was more sensitive to the padding strategy, with only non-hydrophobic padding providing equivalent performance. Notably, the model trained with no padding performs comparably to existing methods, suggesting the padding is not strictly necessary for this task. As with Serrano157, the ESM2 8M embeddings were critical for performance, but increasing the ESM2 model scale had a negative effect. The ablations indicate that the weighted mean and residue padding serve similar functions; removing either results in a minor decrease, whereas removing both causes a significantly larger decline. It is important to note that the residue-level annotations in AmyPro22 are compiled using different experimental approaches and therefore may be of limited accuracy, in particular there may unannotated APRs. Visual inspection of the residue score profiles of Aβ42 and the PB1-F2 residue profiles shows that there is a range of residue scores within the regions labeled as APR, suggesting that finer-grained resolution of residues contributing to aggregation may be possible. Ultimately, these predictions will need to be validated by future experimental work, either through mutations or transplantation of APRs. As PALM can be trained with sequence-level labels, which are readily obtained from experiments, it can continue to improve at APR identification as new data becomes available.

### A combination of limited data and pLM features leads to poor performance when predicting single mutant effects

The ability to predict the impact of single mutations on aggregation is important for guiding the design of new sequences. Given the strong performance of PALM on Serrano157 and NNK4, we assessed whether it could identify single amino acid substitutions in Aβ42 that lead to dominant familial Alzheimer’s disease by increasing rate of aggregation. We found that PALM was unable to identify that these mutations increase aggregation, in part because the residue scores at these positions were already close to the maximum value. Training PALM with NNK1-3, a larger and more diverse dataset than WaltzDB-2.0, resulted in improved performance on this task. The residue score profiles of PALM (NNK1-3) reveal that the values are no longer saturated for the WT sequence, and that the aggregation-inducing mutations consistently increase the scores at multiple positions in the sequence.

Interestingly, we found that training PALM with NNK1-3 and one-hot encodings substantially improved performance on the mutation prediction task over ESM2 8M embeddings. Furthermore, PALM with one-hot encodings also improved over base PALM. Together, these results show that it is both the small amount of data and the ESM2 features that limit performance on this task. We note that CANYA performs remarkably well on the mutational effect task, showing improved AUPRC and enrichment factor over all methods we benchmarked. This suggests the unique features of its architecture: e.g. the self-attention with positional embeddings, may be important for this specific task. Future work can explore the potential of fine-tuning pLMs with larger datasets.

## Conclusion

PALM leverages protein language model embeddings to predict aggregation in peptide sequences equivalent to state-of-the-art existing methods. The architecture predicts residue-level scores which can be used for the identification of aggregation-promoting regions within protein sequences, without being explicitly trained on this task. As new data becomes available, PALM can be improved, as demonstrated by training on the NNK1-3 dataset, which enabled the identification of aggregation-promoting mutations in Aβ42 peptide. We release the code and model weights for PALM, PALM NNK, and PALM NNK OH, so that they may used by the community for aggregation prediction: including screening library for potential amyloid peptides and identifying potential APRs. These models and scientific insights will support the development of therapeutic peptides and the identification of aggregation-promoting mutations relevant to disease.

## CODE AVAILABILITY

Code and model weights are available at https://github.com/novonordisk-research/PALM.

## ACKNOWLEDGEMENTS

We thank Christos Nicolaou, Kay Schaller, Anna Macintyre and other members of Digital Chemistry & Design at Novo Nordisk for their support. We also thank various research teams for publishing open source models, including Burkhard Rost’s group and the ESM2 team. Similarly, we thank the groups who developed AggreProt and ANuPP for their helpful responses to our inquiries.

## Supplementary Information

**Table S1.** A summary of experimental datasets used for model training and evaluation. The curation of the datasets is described in the methods.

| Dataset | Type | Sequence count | Sequence length range | Amyloid (%) | Annotation level |
| --- | --- | --- | --- | --- | --- |
| WaltzDB-padded | Train | 14,148 | [8, 26] | 36.2 | Sequence |
| AmyPro22 | Test | 22 | [35, 660] | 18.3 | Residue (APR) |
| Serrano157 | Test | 158 | [6, 36] | 24.6 | Sequence |
| Aβ42 mutants | Test | 753 | [42] | 1.7 | Sequence |
| NNK1-3 | Train | 100,730 | [1, 20] | 20.7 | Sequence |
| NNK4 | Test | 7,039 | [1, 20] | 23.8 | Sequence |

**Table S2.** Comparison of train and validation loss measured on the WaltzDB-2.0 dataset at the epoch with the lowest recorded validation loss for each ESM2 model size.

| ESM2 size | Binary cross entropy loss |  |
| --- | --- | --- |
|  | Train | Validation |
| 8M | 0.389 | 0.461 |
| 35M | 0.382 | 0.487 |
| 150M | 0.372 | 0.457 |
| 650M | 0.361 | 0.516 |

**Table S3.** Performance comparison between PALM, TANGO, and CANYA (ROC AUC) across Serrano and Amypro datasets segmented by sequence length.

| Model | Serrano157 (ROC AUC) |  |  | AmyPro22 (ROC AUC) |  |  |
| --- | --- | --- | --- | --- | --- | --- |
|  | >18aa | 10–18aa | <10aa | ≤133aa | >133aa | - |
| PALM | 0.85 | 0.92 | 0.90 | 0.67 | 0.69 | - |
| TANGO | 0.77 | 0.94 | 0.99 | 0.61 | 0.66 | - |
| CANYA | 0.81 | 0.85 | 0.80 | - | - | - |

**Table S4.** Spearman correlations between PALM model scores and Kyte-Doolittle hydrophobicity values. *Indicates correlations with p-values above significance cutoff.

| Dataset | Serrano157 | AmyPro22 | NNK4 |
| --- | --- | --- | --- |
| Residue scores vs. hydrophobicity | 0.223 | 0.230 | 0.227 |
| Sequence scores vs. average hydrophobicity | 0.520 | 0.262* | 0.320 |
| Residue class labels vs. hydrophobicity | - | 0.053 | - |
| Sequence class labels vs. average hydrophobicity | 0.412 | - | 0.173 |

**Table S5.** Performance comparison across Serrano, Amypro, and NNK4 datasets, split into two equally sized groups by Kyte-Doolittle hydrophobicity (*↑* = High, *↓* = Low).

| Dataset | Serrano157 |  |  |  |  |  | AmyPro22 |  |  |  |  |  | NNK4 |  |  |  |  |  |
| --- | --- | --- | --- | --- | --- | --- | --- | --- | --- | --- | --- | --- | --- | --- | --- | --- | --- | --- |
| Metric | ROC AUC |  | PR AUC |  | Enrichment |  | ROC AUC |  | PR AUC |  | Enrichment |  | ROC AUC |  | PR AUC |  | Enrichment |  |
| Hydrophobicity | ↑ | ↓ | ↑ | ↓ | ↑ | ↓ | ↑ | ↓ | ↑ | ↓ | ↑ | ↓ | ↑ | ↓ | ↑ | ↓ | ↑ | ↓ |
| PALM | 0.92 | 0.86 | 0.93 | 0.50 | 2.04 | 4.06 | 0.67 | 0.68 | 0.33 | 0.43 | 2.26 | 2.38 | 0.64 | 0.72 | 0.43 | 0.37 | 1.18 | 1.90 |
| PALM NNK | - | - | - | - | - | - | - | - | - | - | - | - | 0.80 | 0.82 | 0.70 | 0.59 | 2.34 | 3.20 |
| PALM NNK OH | - | - | - | - | - | - | - | - | - | - | - | - | 0.75 | 0.75 | 0.57 | 0.48 | 1.89 | 2.71 |
| TANGO | 0.85 | 0.86 | 0.73 | 0.64 | 1.53 | 4.36 | 0.67 | 0.62 | 0.35 | 0.41 | 3.04 | 2.61 | 0.63 | 0.69 | 0.42 | 0.36 | 1.12 | 2.05 |
| Waltz | 0.77 | 0.82 | 0.64 | 0.36 | 1.02 | 2.90 | - | - | - | - | - | - | 0.65 | 0.68 | 0.46 | 0.33 | 1.50 | 1.71 |
| CANYA | 0.83 | 0.74 | 0.80 | 0.29 | 2.04 | 2.90 | - | - | - | - | - | - | 0.79 | 0.80 | 0.66 | 0.57 | 2.17 | 3.08 |
| AggreProt | 0.90 | 0.86 | 0.88 | 0.43 | 1.53 | 2.18 | 0.64 | 0.64 | 0.25 | 0.37 | 2.12 | 2.26 | 0.61 | 0.58 | 0.43 | 0.26 | 1.29 | 1.23 |
| Anupp | 0.86 | 0.78 | 0.87 | 0.30 | 2.04 | 2.90 | 0.64 | 0.67 | 0.31 | 0.40 | 2.61 | 2.45 | - | - | - | - | - | - |
| Aggrescan | 0.76 | 0.75 | 0.84 | 0.19 | 2.04 | 2.90 | 0.62 | 0.68 | 0.30 | 0.40 | 2.50 | 2.40 | - | - | - | - | - | - |
| Pasta | 0.25 | 0.36 | 0.35 | 0.10 | 0.51 | 0.72 | 0.59 | 0.58 | 0.21 | 0.25 | 1.49 | 1.21 | - | - | - | - | - | - |

**Fig. S1.**
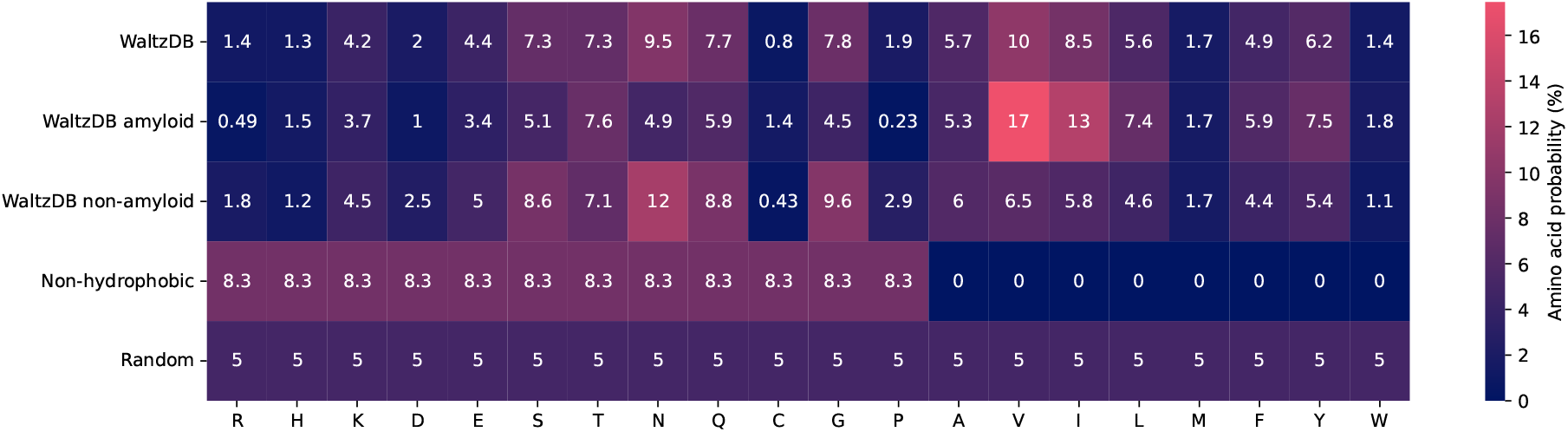
Amino acid probability distributions for datasets used in training and residue padding. Random and non-hydrophobic padding strategies are evenly distributed among subsets of amino acids, while distributions measured on subsets of the WaltzDB dataset are taken from experimentally measured proteins.

**Fig. S2.**
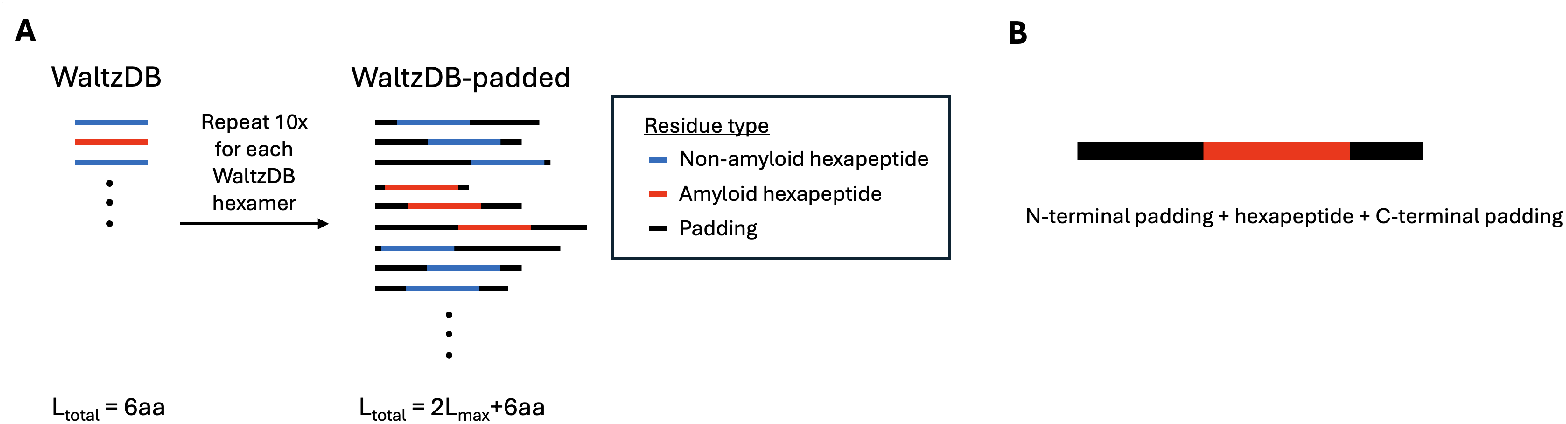
The data augmentation process for WaltzDB. A) A hexapeptide sequence is selected from WaltzDB and then two distinct padding sequences are appended to the beginning and end of the sequence. Amyloid sequences are shown in red, non-amyloid sequences are shown in blue, and padding sequence is shown in black. B) The padded sequences are made up of three distinct substrings: N-terminal padding, the original hexapeptide, and C-terminal padding.

**Fig. S3.**
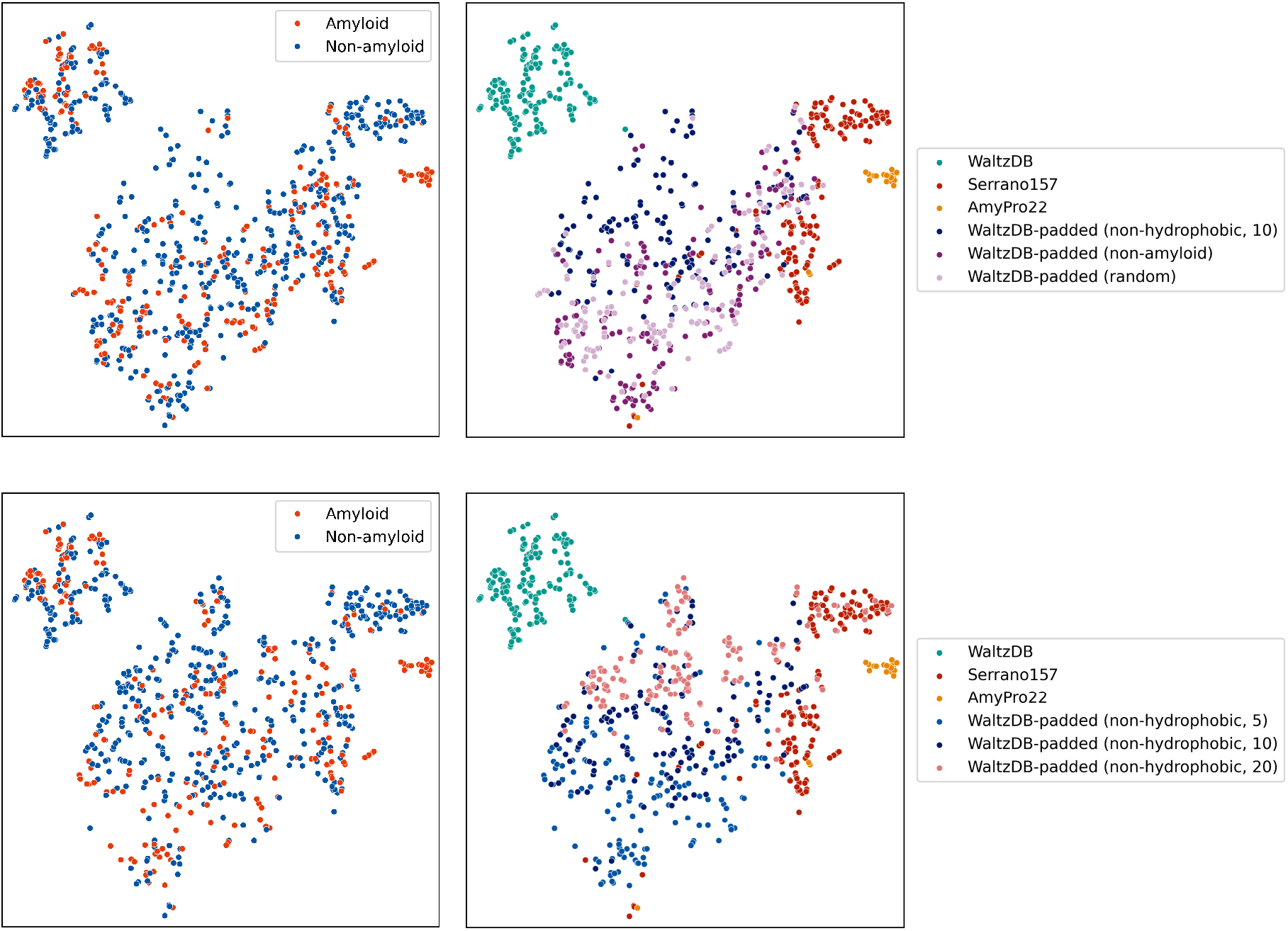
ESM2 embeddings for sequences generated with different padding strategies visualized using t-SNE, colored by label (left column) and dataset (right column). Top row: comparing sequences generated with different residue padding distributions, all with max length 10 aa. Bottom row: comparing sequences generated with the non-hydrophobic residue padding strategy, with different max lengths. Datasets with different maximum padding lengths occupy different regions of embedding space.

**Fig. S4.**
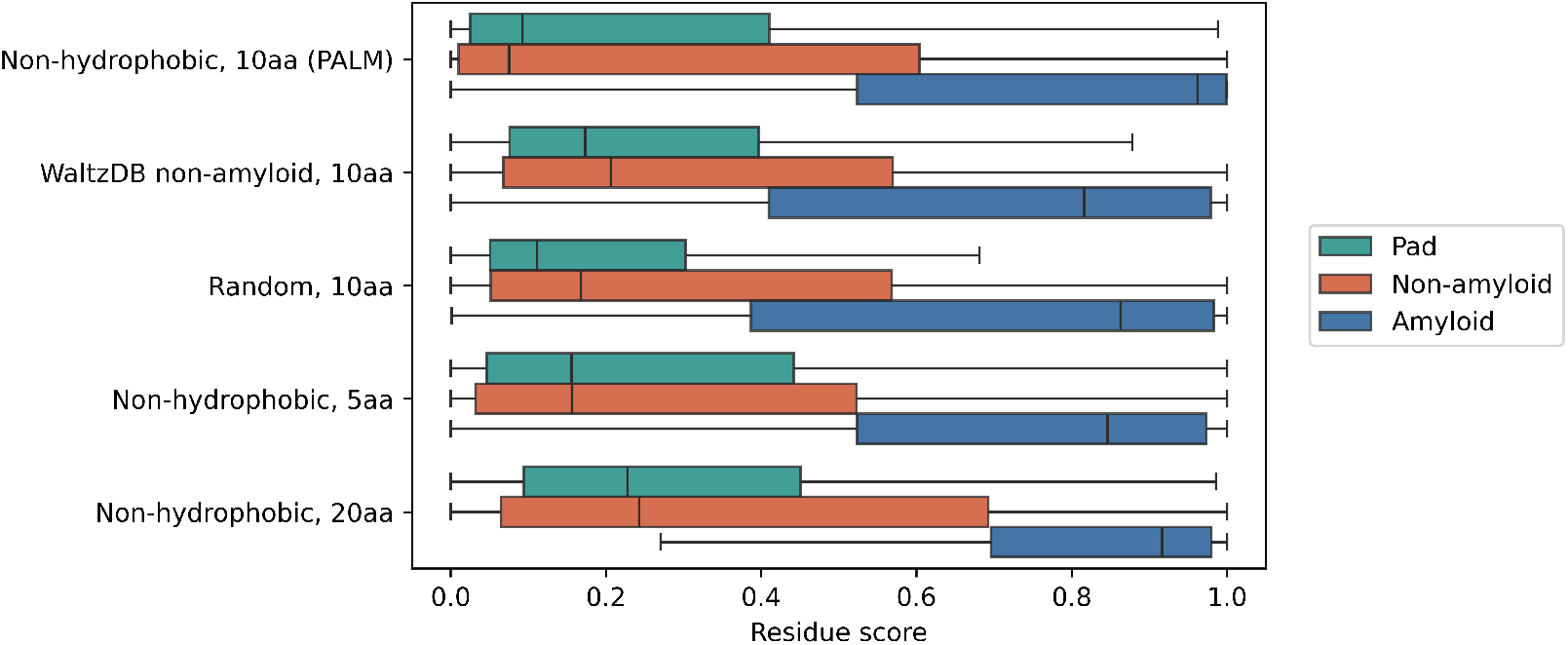
The residue score distributions from models trained on WaltzDB augmented with different padding strategies. The distributions were computed using the validation fold from the WaltzDB training set. Residues are grouped as padding (green), non-amyloid hexapeptides (orange), amyloid hexapeptides (blue).

**Fig. S5.**
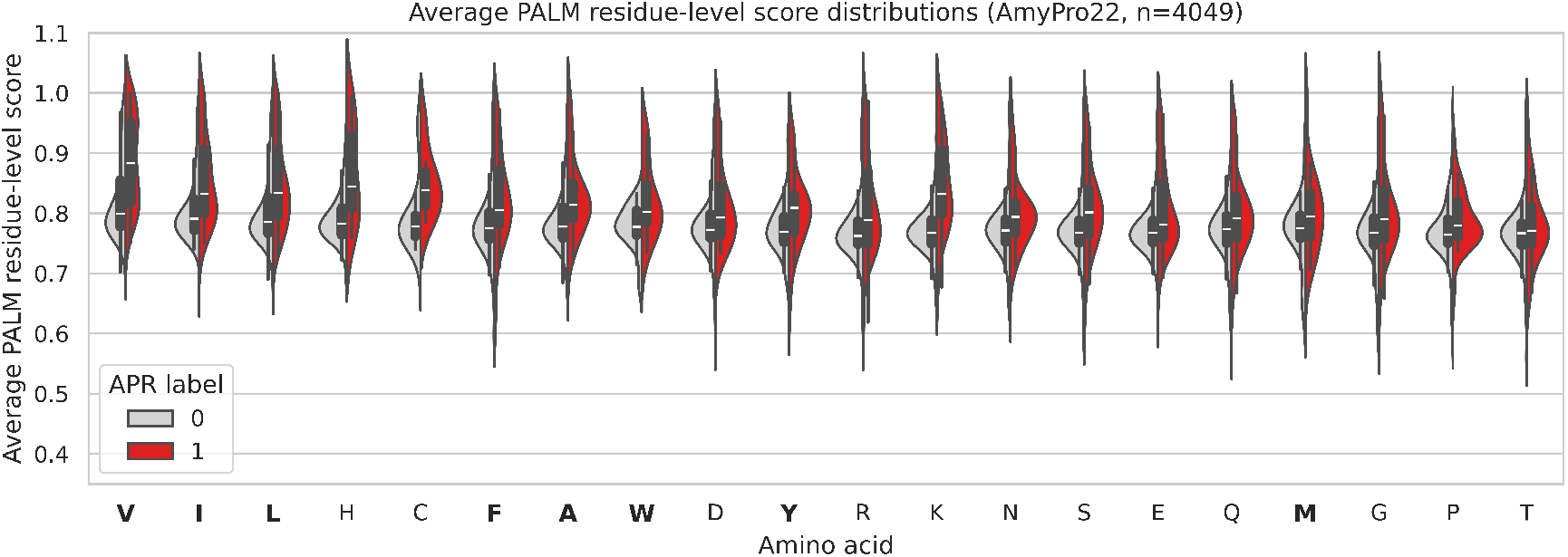
Residue score distributions per amino acid type generated from PALM on the AmyPro22 dataset. Residues not sampled by the non-hydrophobic WaltzDB padding strategy are highlighted in bold.

**Fig. S6.**
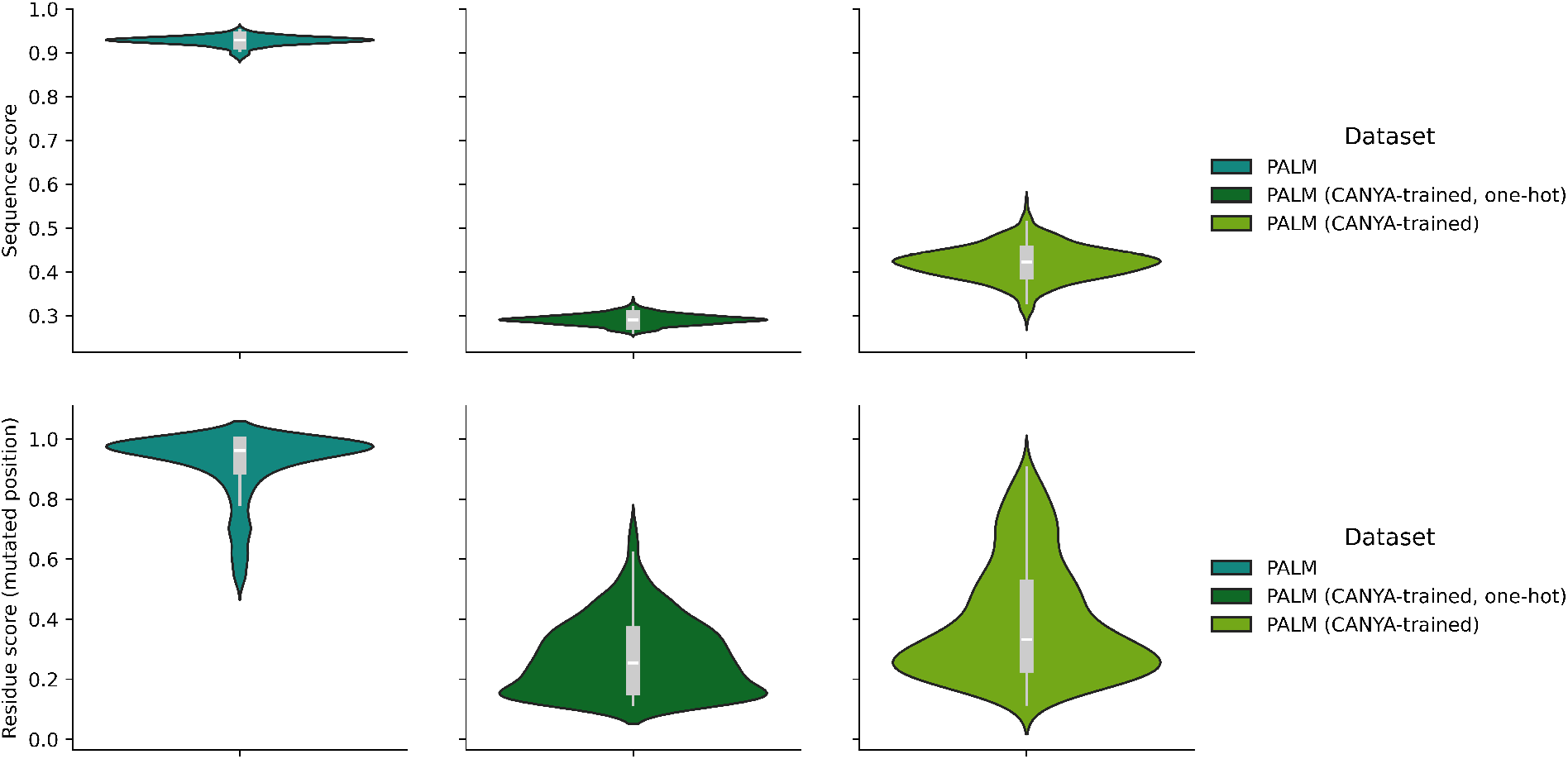
Violin plots of sequence and residue scores taken from the amyloid beta mutant dataset. Scores are taken from three variants of PALM: base, CANYA-trained, and CANYA-trained with one-hot embeddings. Top: sequence scores for all 753 amyloid beta mutants. Bottom: residue scores taken from the mutated position of each sequence.

